# Optimizing the membrane composition of immobilized giant vesicles for effective entrapment and observation of motile bacteria

**DOI:** 10.1101/2025.11.20.689592

**Authors:** Paulo Chu, Liu Yu, Cécile Fradin

## Abstract

Artificial lipid vesicles are an elegant way of producing auto-assembled customizable chambers, and they have been widely studied for entrapping small molecules in the context of drug delivery. However, their use for microorganism entrapment remains largely unexplored. One obstacle is the generation of giant vesicles (GVs) large and rigid enough to contain potentially mobile micron-size organisms in a way that results in efficient cell encapsulation. To address this challenge we developed a modified version of the previously described gel-assisted swelling method, in which immobilized GVs are formed by hydrating a lipid film deposited on agar with a heated solution. We used solutions containing *Magnetospirillum magneticum* strain AMB-1 cells to test for bacterial entrapment. Systematically exploring POPC:DOPS:cholesterol ternary mixtures to vary the surface charge and rigidity of the GVs’ membrane allowed the determination of specific conditions leading to the efficient entrapment of these highly mobile cells. Lipid order measurements with the fluorescent membrane probe Laurdan show that entrapment is associated with a significant decrease in membrane rigidity between the swelling and entrapment steps. The high sensitivity of successful entrapment on DOPS concentration further suggests that membrane fluctuations play a major role in allowing the micron-size bacteria to push through the membrane and break in and out of the GVs.

## 1 INTRODUCTION

Lipid vesicles mediate a wide range of biological functions, from intracellular transport and protein degradation to intercellular communication and waste removal [1, 2]. They are also widely used for bioengineering and biomedical applications, for instance, as extracellular templates in reconstituted membrane studies [3, 4] or as drug delivery systems [5, 6, 7, 8]. What makes them so attractive for these kinds of applications are their excellent biocompatibility, their intrinsic ability to self-assemble, their capacity to encapsulate both hydrophilic and lipophilic molecules, and their unique soft-matter properties. Also important is the fact that these properties can be tailored for each particular application by modifying lipid composition and using additives such as sterols, proteins (ion channels, chemical receptors) or lipopolysaccharides, embedded in or attached to the lipid bilayer.

In this study, we explore the optimal membrane compositions for the entrapment of bacteria in giant vesicles (GVs) with diameters in the range of 1 to 50 *μ*m. The prospect of entrapping bacteria in GVs is attractive for a few reasons. First and foremost, it allows one to monitor the behaviour of individual bacteria in a stable biocompatible environment for extended periods of time, including following the fate of bacterial cultures arising from a single cell [9, 10, 11]. Furthermore, by providing a compartment spacious enough to allow unrestrained motions, GVs allow observing not just cellular growth and division, but also cellular motility. In addition, the behaviour of entrapped and non-entrapped bacteria can be observed in parallel, to compare their response to environmental changes or stimuli applied only to the more easily accessible, non-trapped bacteria. This could be used to directly assess the effect of specific molecules, such as antibiotics, that are unable to cross the membrane of the GVs [10].

The capacity of lipid vesicles to entrap molecules such as fluorescent dyes, therapeutic drugs, proteins, and nucleic acids has been well documented [12, 13, 14, 15, 16]. Molecular entrapment has been shown to depend on variables related to the particles to be entrapped (molecular weight, charge), but also to the vesicles themselves (size, lipid composition, membrane zeta potential), and to the surrounding aqueous solvent (buffer composition, salt concentration, temperature). In contrast, only very little work has been done regarding the entrapment of bacteria in lipid vesicles. In early works, microinjection was used to introduce *Escherichia coli* (*E. coli*) cells in pre-formed giant unilamellar vesicles (GUVs) [17]. More recent studies have showcased instead a droplet-transfer method (also known as inverted emulsion method) to create GUVs already entrapping bacteria [10, 11, 18]. The latter involves the formation of water-in-oil droplets stabilized by a lipid monolayer, which are then transferred by gravity or centrifugation to an aqueous phase while acquiring a second lipid layer, and finally immobilized on a glass substrate. Droplet-transfer allows for efficient and controlled cellular entrapment, but relies on the inner aqueous phase of the vesicle to be denser than that of the surrounding medium, thus requiring a high osmolarity inside the vesicles. It was also shown to yield vesicles with lipid ratios different from those intended [19].

Here we explored the possibility to entrap bacteria using a simpler method to form GVs, known as the swelling method or gel-assisted swelling method, first proposed by Horger *et al*. in 2009 [20, 21]. It is derived from the classic thin-film hydration method (also known as the Bangham’s method) [22], and consists in hydrating a lipid film deposited on a thin partially-dehydrated hydrogel film. Importantly, it allows working with solutions of high ionic strength, essential for bacterial growth [20]. Another advantage is that GVs obtained by gel-assisted swelling are already immobilized on the hydrogel on which they form. We developed a protocol based on this method to entrap a motile strain of bacteria, the Gram-negative facultative anaerobic *Magnetospirilum magneticum* strain AMB-1 (referred to in the following as AMB-1), often used as a model system to study magnetotactic organisms [23, 24]. AMB-1 cells swim thanks to the counter rotation of their helical body and their two flagella, exerting a total propulsive force of *≈*2 pN. [25, 26] We tested a number of lipid mixtures including 1-palmitoyl-2-oleoyl-sn-glycero-3-phosphocholine (POPC), 1,2-di-(9Z-octadecenoyl)-sn-glycero-3-phospho-L-serine (DOPS), and cholesterol, and showed that successful bacterial entrapment using the gel-assisted swelling method critically depends on lipid composition.

## 2 EXPERIMENTAL

### 2.1 Materials

Fluorescent 110 nm-diameter nanobeads (F8803, excitation / emission: 505 / 515 nm) were obtained from Invitrogen (Eugene, Oregon, USA). Laurdan (1-(6-(dimethylamino) naphthalen-2-yl)-dodecan-1-one) was sourced from Cayman Chemical (Ann Arbor, Michigan, USA). 1-palmitoyl-2-oleoyl-glycero-3-phosphocholine (POPC, *T*_*m*_ = −2°*C*), 1,2-dioleoyl-sn-glycero-3-phospho-L-serine (sodium salt) (DOPS, *T*_*m*_ = −11°*C*), and cholesterol (ovine wool, 98%; *T*_*m*_ = 141°*C*) were purchased from Avanti Polar Lipids (Alabaster, Alabama, USA). Alexa Fluor 488 was obtained from Invitrogen (Carlsbad, CA, USA).^1^

### 2.2 Cell culture

*Magnetospirillum magneticum* strain AMB-1 was obtained from the American Type Culture Collection (ATCC®, catalog number 700264). Cells were cultured in a growth culture medium originally described for the species MSR-1 in Le Nagard *et al*. [27]. This medium, referred to below as MSGM-LN (MagnetoSpirillum Growth Medium Le Nagard) contains 10mM HEPES, 16 mM KC3H5O3, 4 mM NaNO3, 0.7 mM KH2PO4, 0.6 mM MgSO4.7H2O, 0.05 mM ferric citrate, trace mineral supplements (diluted 1000 time from the stock provided by ATCC), 3 g/l soybean peptone, 0.1 g/l yeast extract, and has a pH adjusted to 7. All AMB-1 populations are racetracked before sub-culturing, a method to retain magnetotacticity.[28]

### 2.3 Formation of bacteria-entrapping GVs

GVs were formed following the gel-assisted swelling method [20], with small adjustments to adapt it for bacterial entrapment. ^2^

#### Mixed lipid and agarose film formation

A chloroform solution with the desired lipids and cholesterol molar ratios (total molarity 2.86 mM) was prepared in a glass vial from stock solutions of POPC, DOPS, and cholesterol (each in chloroform). Laurdan was added to the aliquots of these mixtures at a 1:100 molar ratio [29] for fluorescence imaging. This operation was performed in a ductless fume hood, using a glass syringe extensively cleaned with chloroform. The headspace of the glass vial was purged with argon, the vial capped, sealed with parafilm, and the solution stored at −20°*C*.

A double deionized water (17 MΩ/cm) solution with 2% w/w Agar-A was alternatingly microwaved for 5 s and swirled, until the agar was fully dissolved. Using tweezers, a 24×60 mm #1.5 microscope coverslip was briefly placed to float on top of the hot agar solution, after which it was picked up and excess agar allowed to drip for a few seconds. The coverslip was then placed on a ≈ 50°*C* heating block, agar side up, protected from dust and debris with a lid, and given 45 min to partially dehydrate. The thin agar layer is then translucent to transparent.

Working in a ductless fume hood, a clean glass syringe was used to deposit 12.5 µL of the lipid solution in small droplets at one edge of the agar layer. The syringe needle was then laid flat on the surface and used to spread these droplets across the agar surface. This operation was repeated at the opposite edge of the agar layer. The agar- and lipid-coated coverslip was then left for 15 min at room temperature in the fume hood to let the chloroform to evaporate. Two thin parafilm strips were placed on the coverslip less than 18 mm apart, after which an 18×18 mm #1.5 coverslip pre-heated to 60°*C* was bridged atop the parafilm strips. A light push on the top coverslip with tweezers allowed binding the two glass coverslips together with the parafilm, creating a *≈* 50 µm thick chamber between them. The edges of that chamber were partially sealed with wax, leaving one small opening on each non-parafilm side diagonally.^3^

#### Lipid film hydration and bacterial entrapment

Different hydrating solutions were used as needed: Dulbecco’s Phosphate Buffered Saline (DPBS), DPBS containing fluorescent nanobeads (at a concentration of 5.7 beads/µm^3^ or approximately 10 nM), DPBS containing 5 *μ*M of Alexa Fluor 488, or a bacterial culture (AMB-1 grown in MSGM-LN). The solution was pre-heated in a water bath at 40°*C*, then 30 *μ*L was pipetted into one of the openings between the two coverslips, so as to completely fill up the chamber, and the two small openings sealed with wax to prevent evaporation. The sample was then left to equilibrate back to room temperature.

### 2.4 Light microscopy

A Nikon TS100 microscope with a Plan-Apochromat 60x/0.95 NA air objective and a mounted high-speed Allied Vision GigE 680 camera was used for transmitted light microscopy imaging. For confocal imaging, a Zeiss LSM 980 Inverted Confocal with a Plan-Apochromat 63x/1.40 NA oil or a Plan-Apochromat 20x/0.8 air objective and a GaAsP-PMT was used. Fluorescence was excited with a 405 nm laser line (blue/cyan emission channel) for Laurdan and a 488 nm laser line (green emission channel) for Alexa Fluor 488 and fluorescent nanobeads.

### 2.5 Characterization of lipid order

Lipid order was assessed using the membrane dye Laurdan, whose fluorescence emission is sensitive to the polarity of its local environment [30].

#### Vesicle preparation

Simple gentle hydration was used to form vesicles for use in fluorescence spectroscopy experiments. Chloroform solutions with the desired POPC, DOPS, and cholesterol molar ratios were prepared as explained above. Laurdan was added to the lipid mixtures at a 1:100 molar ratio [29]. The final molar concentration was 2.41 mM. A 30 *μ*L volume of this solution was placed in a 2 mL glass vial, and the chloroform allowed to evaporate in a ductless fumehood. The dried lipid mixture was then hydrated by pipetting up and down 200 *μ*L of ΔMSGM-LN (MSGM-LN lacking yeast and soytone extracts, ingredients found to be slightly fluorescent when excited at 365 nm). Vesicle formation was confirmed by the solution becoming cloudy.

#### Fluorescence spectroscopy measurements

A Tecan Safire^2^ microplate reader was used for fluorescence spectroscopy measurements. 100 *μ*L of each vesicle solution was dispensed into a 96-well plate with transparent polystyrene-bottom (Thermo Fischer Scientific, catalog number 165305). Laurdan was excited at 365 nm, and its emission spectrum measured between 380 nm and 600 nm (emission and excitation bandwidths: 2.5 nm; emission wavelength step size: 1 nm). An integration time of 40 *μ*s was used and the manual gain was set to 100. The emission spectrum of ΔMSGM-LN was recorded at each temperature, and subtracted from the emission spectra recorded from vesicle solutions.

#### Estimate of the generalized polarization parameter

When in a lipid membrane, Laurdan has two fluorescence emission peaks, around 440 nm and 490 nm, corresponding to Laurdan molecules found either in a lipid ordered or lipid disordered environment, respectively. The relative amplitude of these two peaks is thus an indication of membrane order. This balance is captured in a dimensionless quantity known as generalized polarization (GP), defined as [29, 30]:

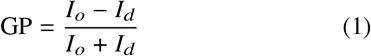

In the above equation, *I*_*o*_ and *I*_*d*_ are the intensities of the Laurdan fluorescence emission at the position of the peaks found around 440 and 490 nm, respectively. The GP ranges from −1 to 1 and its value is directly correlated to membrane order.

To obtain the GP of a specific vesicle population, the emission spectrum (after subtraction of the ΔMSGM-LN emission spectrum) was fitted to a double log-normal function, as suggested by Bacalum *et al*. [31]:

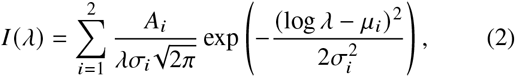

where *A*_*i*_, *μ*_*i*_ and *σ*_*i*_ correspond to the amplitude, position, and width of each of the two emission peaks. The peaks positions were found to be *μ*_1_ *≈* 434 nm and *μ*_2_ *≈* 487 nm. The GP was obtained by calculating the intensities *I*_*o*_ = *I* (*μ*_1_) and *I*_*d*_ = *I* (*μ*_2_) from Eq.2 using the fit parameters, then inserting these values into Eq.1.

## 3 RESULTS AND DISCUSSION

### 3.1 Particle entrapment in GVs formed by gel-assisted swelling

To entrap live motile bacteria (in this case AMB-1) in immobilized GVs for long periods of time, we looked for a GV formation method that would produce already immobilized GVs in bacterial growth media with potentially high ionic strength. We chose to use the gel-assisted swelling method, [20, 21] in which a partially dehydrated agarose film is coated with a lipid film on a glass surface, then hydrated with a buffer solution. This method was specifically developed as an easy way to form GVs regardless of the ionic strength of the hydrating solution [20]. In the present work, the hydrating solution contained particles to be entrapped in the GVs, most often live AMB-1 cells in culture medium (as illustrated in Fig. 1). We chose to use lipid mixtures made of three components: POPC (to form GV efficiently [20]), cholesterol (to give higher mechanical resistance to the membrane), and DOPS (to obtain a negatively charged GV membrane). The one notable change made to the original protocol, is that the hydrating solution was pre-heated to a temperature of 40°*C*.

**Figure 1:**
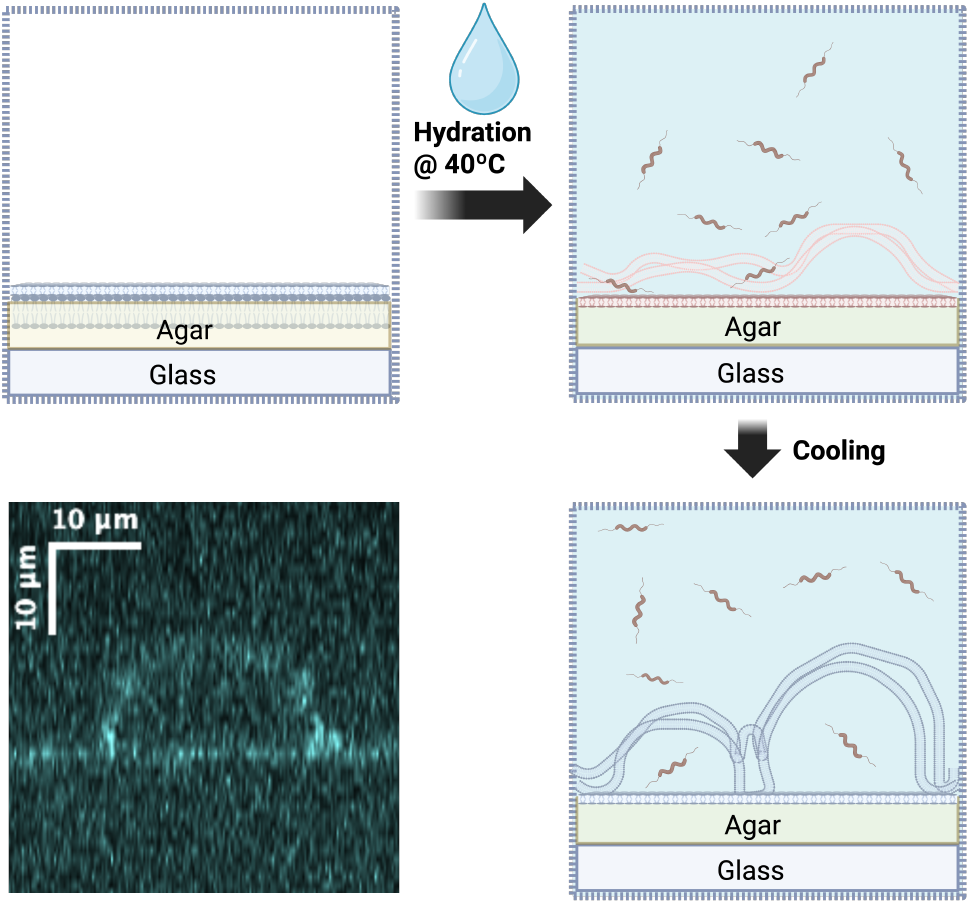
Principle of bacterial entrapment in GVs formed by gel-assisted swelling. Three steps are involved in the formation of GVs entrapping bacteria: deposition of a lipid film on top of a thin agar layer, hydration with a warm aqueous bacterial culture, and finally cooling to room temperature during which the swelling occurs. A vertical confocal image of a formed GV immobilized on the agar layer is shown at the bottom-left. Figure created with BioRender.com/p3txqfs.

To verify whether GVs formed by gel-assisted swelling were able to entrap small particles, we first performed preliminary experiments where the mixed agarose-lipid films were hydrated with DPBS solutions containing either the fluorescent dye Alexa Fluor 488, or fluorescent nanobeads. Confocal fluorescence microscopy was then used to image the GVs and assess the success of fluorophore entrapment (Fig. 2). We observed that some combinations of lipid compositions led to dye entrapment, while others did not, emphasizing the importance of optimizing experimental parameters, and in particular lipid composition, for each type of particles being entrapped. None of the tested compositions entrapped any fluorescent nanobeads, illustrating the difficulty for inert colloidal particles to penetrate the forming GVs.

**Figure 2:**
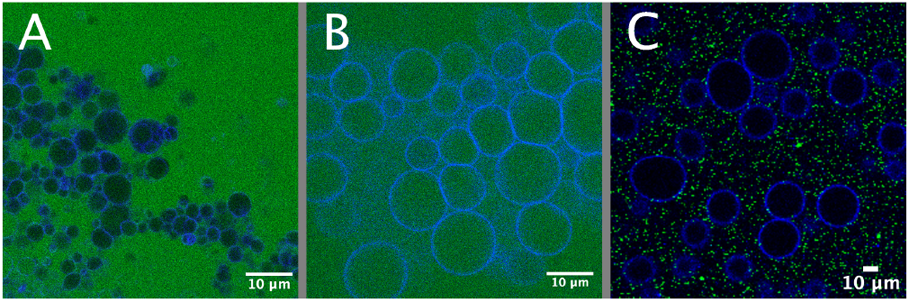
Gel-assisted swelling leads to particle entrapment only in certain conditions. Confocal microscopy images of GVs labeled with Laurdan (blue) formed by gel-assisted swelling using either (A,B) a DPBS solution with 5 *μ*M of Alexa Fluor 488 (green) or (C) a DPBS solution with 10 nM 110 nm-diameter fluorescent nanobeads (green). The lipid composition used was 40:60 (POPC:cholesterol) in (A) and 60:20:20 (POPC:DOPS:cholesterol) in (B) and (C).

**Figure 3:**
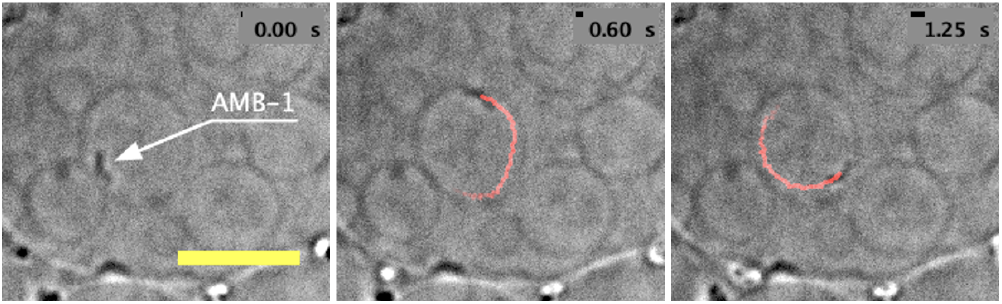
Example of AMB-1 cell entrapped in a GV formed by gel-assisted swelling. Time-lapse series of light microscopy images showing an entrapped AMB-1 cell in a GV with lipid composition of 60:25:15 (POPC:DOPS:cholesterol). In each panel, the cell trajectory for the last 0.55 seconds is shown in red. Scale bar is 10 *μ*m.

In order to achieve bacterial entrapment, the hydrating solution was replaced by an AMB-1 culture in MSGM-LN medium, harvested 3 days after inoculation (OD *≈* 0.24 at 633 nm). Then the formed GVs were imaged with transmitted light microscopy, as the agar layer on top of the coverslip is thin enough to allow good-quality optical imaging and the cells are large enough to be detected without any fluorescence labeling. The motility of AMB-1 cells made it easy to confirm entrapment, since entrapped cells could be clearly identified by their constrained circular trajectories, as shown in Fig.3.^4^ This particular example shows that gel-assisted swelling can achieve bacterial entrapment. However, as in the case of the dye, we observed that entrapment was not observed in all conditions. In particular, focusing first on POPC:cholesterol compositions, we found that using a room temperature solution resulted in GV formation but not in bacterial entrapment. This led us to systematically pre-heat the hydrating solution to 40°*C*. This particular temperature was chosen as higher temperatures are detrimental to the health of AMB-1 cells.

These preliminary experiments confirmed that gel-assisted swelling can lead to bacterial entrapment, with the advantage of frugality and speed. The only required ingredients are the agar and the lipid constituents (in the present case, POPC, DOPS, and cholesterol). The primary role of the agar layer and of its irregular topology is to provide nucleation sites for the GVs, ensuring robust formation for a large range of lipid and buffer compositions [32, 20]. However, as an added benefit, the interaction between the agar and the lipid film also ensures that the formed GVs remain pinned to the substrate. When captured in confocal images, most GVs appear as immobilized hemispherical compartments solidly anchored to the agar layer, ideal for the observation of entrapped particles (Fig. 1). Immobilization of already formed GVs is of course possible, for example using biotin-neutravidin interactions to bind them to a surface [10, 33] or by trapping them in a hydrogel [34] or in a microfluidic chip [11]. However, this extra step might introduce additional challenges when the GVs are entrapping live cells. For example, vesicle immobilization inside an agarose gel requires adding the pre-formed vesicles to liquid agarose at around 62°*C* [34], making it incompatible with the encapsulation of living cells or temperature-sensitive molecules.

### 3.2 Successful entrapment of AMB-1 in GVs formed by gel-assisted swelling critically depends on lipid composition

To make the entrapment of AMB-1 cells more robust, we looked into the influence of lipid composition, as in the case of bacterial entrapment the composition of the hydrating medium and the accessible temperature range are somewhat constrained. We systematically explored a large range of possible POPC:DOPS:cholesterol ternary mixtures, keeping the rest of the protocol unchanged. For each studied lipid composition, we evaluated whether GVs were formed and whether AMB-1 bacteria were entrapped in the GVs. Observation of the GVs formed with each lipid mixture was done using light microscopy between 5 and 30 min after sample hydration. Entrapment efficiency was estimated by recording a video with on average 5 (and between 2 to 7) unique randomly selected areas (211 × 211 *μ*m each) in each sample. The number of live entrapped bacteria was then counted and divided by the total imaged surface area, giving an estimate of entrapment density. We deemed that successful entrapment was achieved when the entrapment density was greater than 20 AMB-1 cells per mm^2^. The result of this systematic exploration is summarized in the ternary diagram shown in Fig. 4. As expected, we found that GVs readily form for almost all lipid compositions, with the notable exception of some DOPS:cholesterol only mixtures. This is in agreement with the previous finding that gel-assisted swelling successfully leads to GV formation for a wide range of lipid compositions [20]. In contrast, successful bacterial entrapment was observed only for a narrow range of lipid mixtures, found along what seems like an almost linear path within the lipid mix ternary diagram (green dots in Fig.3). The most reliable bacterial entrapment conditions seemed to center around 60:20:20 (POPC:DOPS:cholesterol), where up to *≈* 200 cells/mm^2^ were found to be entrapped. Favorable conditions are also found at higher cholesterol concentrations, but entrapment success is not as consistent.

**Figure 4:**
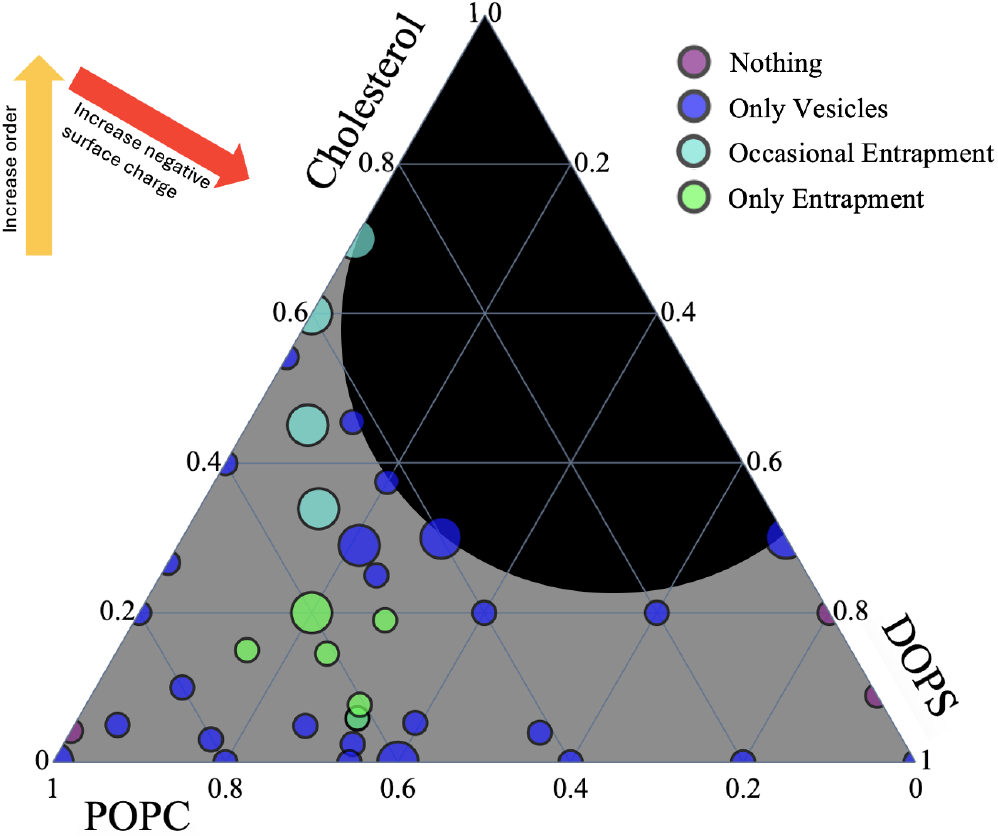
Ternary diagram showing which molar ratios of phospholipid (POPC, DOPS) and cholesterol lead to successful GV formation and bacterial entrapment. Different POPC:DOPS:cholesterol mixtures were used to entrap AMB-1 cells in GVs formed by gel-assisted swelling, and successful GV formation and bacterial entrapment were assessed by light microscopy. Compositions supporting successful AMB-1 entrapment in GVs are indicated by green dots, those for which successful entrapment is observed in some samples but not others by cyan dots, those leading to GV formation but not to AMB-1 entrapment by blue dots, and those not leading to GV formation by purple dots. Larger markers identify compositions that have been assessed repeatedly. The black area is not accessible as it is above the cholesterol saturation limit.

**Figure 5:**
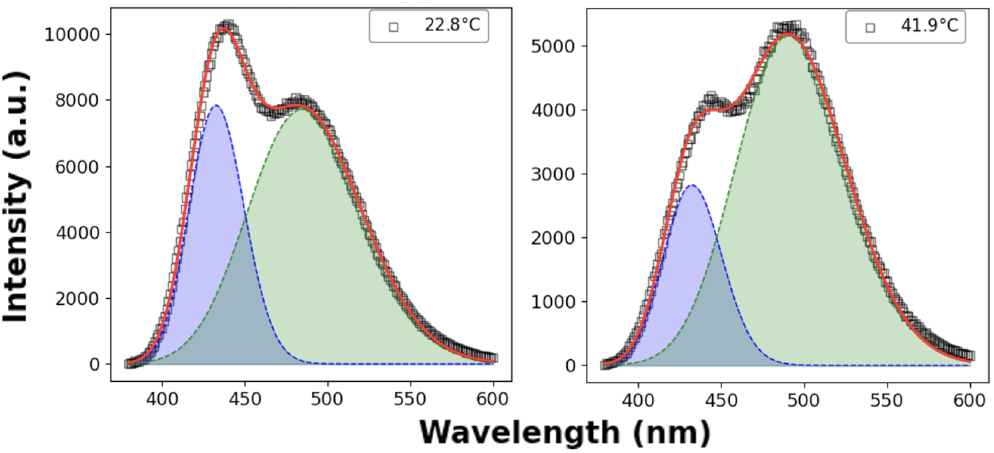
Emission spectrum of Laurdan in POPC vesicles at 22.8°*C* (left panel) and 41.9°*C* (right panel), showing two emission peaks and a clear change in their relative amplitude upon temperature increase. The red line shows the fit of the data with Eq. 2. The blue and green shaded areas show the contributions of the emission peaks corresponding to ordered and disordered lipid environments, respectively.

Our data highlights the importance of all three membrane components (POPC, DOPS, and cholesterol) in order to achieve bacterial entrapment. For mixtures containing only phospholipids, i.e. various fractions of POPC and DOPS and no cholesterol, while GVs are readily forming, they do not support AMB-1 entrapment. When POPC is the only phospholipid in the mix, cholesterol needs to be added at a very high concentration of 60 mol% or higher before any entrapment is observed - and even then, entrapment is not observed reproducibly. This is close or above the known 66 mol% solubility limit of cholesterol in POPC membranes [35]. In these conditions cholesterol crystals might be present and possibly make it easier for the bacteria to penetrate the forming GVs. The formation of cholesterol crystals would also mean that the lipid composition of the formed GVs no longer faithfully reflects the intended cholesterol molar ratio, maybe explaining in part the inconsistency of the results obtained for these very high cholesterol content mixtures.

The addition of a small amount of DOPS allows more robust and consistent entrapment, and at lower cholesterol concentration. Up to a point, cholesterol and DOPS seem to compensate for each other: The more DOPS is added to the membrane of the GVs, the less cholesterol is needed to achieve entrapment - at 30 mol% DOPS, as little as 10 mol% cholesterol is enough to produce robust entrapment. The headgroup of DOPS carries a negative charge at neutral pH due to its serine moiety, thus an increase in DOPS concentration translates into an increase in the surface charge of the GVs. An increase in surface charge generally increases membrane bending rigidity [36], as adding cholesterol does [37]. This suggests that the compensation effect observed between DOPS and cholesterol is the result of entrapment occurring at a precise membrane rigidity (achievable by addition of either molecule) - low enough to allow AMB-1 penetration in the GVs while they are forming at high temperature, but high enough to prevent their escape later at room temperature. A recent experimental study has shown that micron-sized beads could successfully break through the membrane of GUVs if pushed with a force on the order of a few pNs (e.g. on the order of the propulsive force of bacteria), with the minimal force required depending on the bending rigidity of the membrane.[38] Thus in entrapping conditions, the increase in temperature during the hydration step of the entrapment protocol may decrease the rigidity of the GUVs just enough to make it temporarily possible for AMB-1 cells to penetrate GUVs.

We also note that entrapment was never observed above 35 mol% DOPS. AMB-1 is a gram-negative bacteria that belongs to the alphaproteobacteria class [39, 40]. Bacterial species in that class have large quantities of phosphatidylglycerol (PG) and cardiolipins (CL) in their inner and outer membranes, both negatively charged phospholipids [41]. Thus the membrane of AMB-1 cell has a negative surface charge, a common feature of gram-negative bacteria. Large amount of DOPS might keep the AMB-1 cells away from the lipid membrane, never giving them the chance to break into forming or existing GVs.

### 3.3 Fluidity of lipid membranes supporting AMB-1 entrapment

To explore the hypothesis that successful entrapment of AMB-1 cells is connected to the mechanical rigidity of the GVs, we quantified lipid ordering using the fluorescence emission properties of the lipophilic membrane dye Laurdan, which strongly depend on local lipid arrangement. [29, 42] We prepared suspended vesicles with different POPC:DOPS:cholesterol compositions in a version of the AMB-1 culture medium (ΔMSGM-LN) lacking yeast and soytone extracts, since these two ingredients were found to introduce a fluorescence background. The vesicles were formed by gentle hydration of a lipid film containing 1 mol% Laurdan. The Laurdan fluorescence emission spectrum was recorded for temperatures between 22.5 and 42.5°*C* covering the processing range encountered during entrapment experiments. All spectra exhibited, as expected, two clear Laurdan emission peaks, as illustrated in Fig.5. The relative amplitude of these fitted peaks was used to calculate the GP coefficient which is directly related to membrane ordering, as detailed in the method section.

We measured the GP of the lipid mixtures along two specific axes in the ternary lipid diagram - for binary POPC:cholesterol mixtures, and for ternary POPC:DOPS:cholesterol mixtures with equal cholesterol and DOPS amounts, each of them crossing the region in which AMB-1 entrapment is obtained (Fig. 4). For all studied lipid compositions, the GP gradually decreases as temperature increases (Fig. 6). This gentle decrease confirms that the membranes remain in the same liquid state in the considered temperature range, as expected since they are all above their melt transition temperature, *T*_*m*_. It also shows that they become progressively less ordered as the temperature increases. This confirms that the step added to the protocol to achieve entrapment, namely the pre-heating of the hydrating solution to 40°*C*, results in a temporary increase in membrane fluidity by about 0.3 GP when compared to room temperature. This higher fluidity is what might allow bacteria to enter forming GVs.

**Figure 6:**
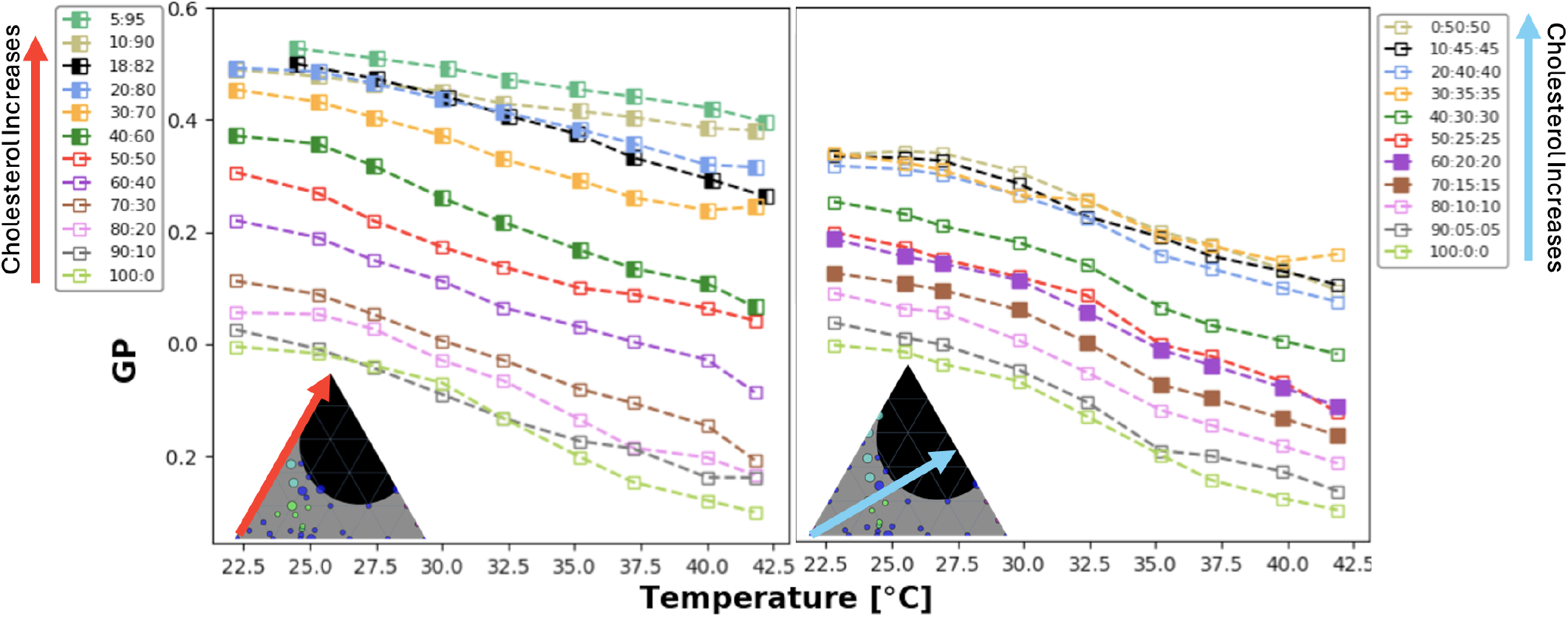
Fluidity of binary and ternary phospholipid and cholesterol mixtures. GP of various mixtures along two different axes in the ternary diagram shown in Fig. 4 as a function of temperature between 22.5 and 42.5°*C*. Filled-in markers indicate a lipid mixture supporting AMB-1 entrapment and half filled-in markers indicate mixtures that have mixed results of vesicles and entrapment. (Left) GP values for binary POPC:cholesterol mixtures. (Right) GP values for POPC:DOPS:cholesterol mixtures with equal molar amounts of cholesterol and DOPS.

Along both the axes that were explored, we observe an increase in membrane order (increase in GP) as more cholesterol is added (Fig. 6), as expected from previous studies [43, 44]. Above a certain amount of cholesterol, the value of the GP saturates, probably reflecting the fact that the limit of solubility for cholesterol has been reached. As expected, this limit depends both on lipid composition and on temperature. In (POPC:cholesterol) membranes, saturation is observed around 60-70 mol% at low temperature (consistent with previously reported measurements of cholesterol solubility in POPC membranes [35, 45]), and around 90-95 mol% at high temperature. In (POPC:DOPS:cholesterol) membranes the saturation limit is significantly lower (also consistent with previous measurements showing that cholesterol is less soluble in membranes made of phospholipids with PS headgroups than in those made of phospholipids with PC headgroups [45]), from 35-40 mol% at low temperature to around 45 mol% at high temperatures.

Our data also shows that an increase in DOPS results in a decrease in membrane fluidity: If one compares the 80:20 (POPC:cholesterol) curve (Fig. 6, left panel, empty pink symbols) to the 60:20:20 (POPC:DOPS:cholesterol) curve (Fig. 6, right panel, filled purple symbols), which differ by a 20 mol% increase in DOPS, a clear 0.2 GP increase is observed across the whole temperature range. This confirms the hypothesis that cholesterol and DOPS can compensate for each other when looking to increase membrane fluidity. However, when specifically considering membranes capable of entrapping AMB-1 cells (denoted by filled symbols in Fig. 6), there is no direct correspondence between the GP of POPC:cholesterol and POPC:DOPS:cholesterol entrapping lipid compositions - the POPC:cholesterol entrapping membranes have higher GPs than the POPC:DOPS:cholesterol entrapping membranes. Thus GP on its own is not a good predictor of successful bacterial entrapment in GVs.

The fact that the optimal Laurdan GP range for bacterial entrapment differs when membrane rigidification is achieved via addition of DOPS versus cholesterol can be reconciled with the idea that successful trapping correlates with membrane rigidity, if we examine what GP is sensitive to. In effect, GP reports on the possibility of rapid dipolar relaxation and local penetration of water molecules in the membrane, and thus on lipid packing (i.e. membrane fluidity) rather than bending stiffness (i.e. membrane rigidity) [43, 46, 44]. Insertion of cholesterol into the bilayer increases acyl-chain order and hence directly decreases fluidity (only secondarily increasing bending stiffness), an effect that Laurdan GP will readily capture. By contrast, charged lipids such as DOPS rigidify the membrane by suppressing small-scale thermal undulations (increasing bending stiffness) [47]. But they might not decrease lateral fluidity to the same extent (especially when the charged lipid is unsaturated, as is the case for DOPS, with a double bond in each of its tails), so their effect might not influence GP as strongly [48]. Local phospholipid packing is known to influence the permeability of membranes to small molecules and proteins, and therefore their encapsulation in vesicles [49]. On the other hand, trapping of larger particles is more likely to be influenced by membrane binding stiffness. Indeed, a correlation has been predicted and experimentally demonstrated between the uptake of micron-size particles into GUVs and the presence of membrane undulations, whose amplitude decreases with membrane rigidity [38]. Our observation that only a small amount (15-20 mol%) of DOPS is sufficient to achieve optimal entrapment is thus consistent with the idea that the thermal fluctuations of the membrane play a major role in allowing bacteria to force their way into GVs - and their absence in preventing them from escaping.

## 4 CONCLUSIONS

We demonstrated the successful entrapment of live bacteria in GVs formed by gel-assisted hydration and swelling. This method, originally developed for the robust formation of GVs [20], proved well-adapted for motile bacteria entrapment - with the only adjustment made an increase in the temperature of the solution used to hydrate the lipid film. Gel-assisted swelling allows the formation of immobilized GVs entrapping live bacteria in physiologically relevant conditions (pH, osmolarity, temperature) and without any accessory molecules (e.g. neutravidin and biotin, glucose, or density gradients).

Whereas gel-assisted GV formation is very robust, bacterial entrapment is extremely sensitive to lipid composition. A ternary lipid mixture proved necessary for optimal entrapment, where POPC was supplemented with small amounts of cholesterol and DOPS. Based on our observations, we speculate that entrapment relies on bacteria pushing their way (using their *≈* pN propulsive force) through the relatively low rigidity membrane of GVs forming around 40°*C*, and on later being unable to escape due to an increase in membrane rigidity at lower temperature. As cholesterol and DOPS provide different ways of achieving membrane rigidity, they can only partially compensate for each other, and DOPS more efficiently suppresses the membrane fluctuations that govern micron-size particle uptake into GVs. Additionally, the negative charge of DOPS might contribute to preventing interactions between the negatively charged bacterial membrane and the GV membrane.

In conclusion, the modified gel-assisted swelling method described in this study presents many useful characteristics when it comes to the entrapment of live mobile bacteria, including simplicity, biocompatibility, and the potential to customize membrane constituents and additives. It might, however, require lipid composition optimization to be adapted to the entrapment of different bacterial species. The entrapping GVs formed using this method should be useful in fields ranging from particle tracking to biochemical microreactor studies and artificial cell models[50].

## Supporting information

Supplemental Figure 1, Table 1, Video 1 description

## AUTHOR CONTRIBUTIONS

P. C.: conceptualization, methodology, investigation, visualization, writing. L. Y.: conceptualization, methodology. C. F.: conceptualization, methodology, funding acquisition, writing.

## ACKNOWLEDGMENTS

We are grateful to Dr. Alex Adronov and Dr. Qiyin Fang for the use of equipment in their laboratories. Confocal imaging was performed at the Center for Advanced Light Microscopy (CALM) at McMaster. This research was funded by the Natural Sciences and Engineering Research Council of Canada (NSERC) through a discovery grant to C.F. (RGPIN-2025-07048).

## SUPPLEMENTARY MATERIAL

Supplementary information is available. The supplementary information includes 1 figure, 1 movie, and 1 table.

## Footnotes

1 A more detailed list of materials and their sources, in particular the ingredients necessary to prepare the growth medium described below, is provided in Supplementary Table 1.

2 A detailed illustration of the experimental protocol is provided in Supplementary Figure 1.

3 See Supplementary Material and Supplementary Fig. 1 for more details.

4 Full video is available as supporting information (Supplementary Video 1).

## REFERENCES

[1] A Longatti and SA Tooze. “Vesicular trafficking and autophagosome formation”. In: Cell Death & Differentiation 16.7 (2009), pp. 956–965.

[2] Guillaume Van Niel, Gisela d’Angelo, and Graça Raposo. “Shedding light on the cell biology of extracellular vesicles”. In: Nature reviews Molecular cell biology 19.4 (2018), pp. 213–228.

[3] Hai Shen, Trevor Lithgow, and Leann L. Martin. “Reconstitution of membrane proteins into model membranes: Seeking better ways to retain protein activities”. In: International Journal of Molecular Sciences 17.3 (2016), p. 326. doi: 10.3390/ijms17030326.

[4] Ralf P. Richter, Rémi Berat, and Alain R. Brisson. “Formation of solid-supported lipid bilayers: An integrated view”. In: Langmuir 22.8 (2006), pp. 3497–3505. doi:10.1021/la052687c.

[5] Theresa M. Allen and Pieter R. Cullis. “Liposomal drug delivery systems: From concept to clinical applications”. In: Advanced Drug Delivery Reviews 65.1 (2013), pp. 36–48. doi: 10.1016/j.addr.2012.09.037.

[6] Giuseppina Bozzuto and Agnese Molinari. “Liposomes as nanomedical devices”. In: International Journal of Nanomedicine 10 (2015), pp. 975–999. doi: 10.2147/IJN.S68861.

[7] Latifa W. Allahou, Seyed Yazdan Madani, and Alexander Seifalian. “Investigating the Application of Liposomes as Drug Delivery Systems for the Diagnosis and Treatment of Cancer”. In: International Journal of Biomaterials (2021), p. 3041969. doi: 10.1155/2021/3041969.

[8] Antonio José Guillot et al. “Skin Drug Delivery Using Lipid Vesicles: A Starting Guideline for Their Development”. In: Journal of Controlled Release 355 (2023), pp. 624–654. doi: 10.1016/j.jconrel.2023.02.006.

[9] ID Vladescu et al. “Filling an emulsion drop with motile bacteria”. In: Physical review letters 113.26 (2014), p. 268101.

[10] Masamune Morita, Kaoru Katoh, and Naohiro Noda. “Direct observation of bacterial growth in giant unilamellar vesicles: a novel tool for bacterial cultures”. In: ChemistryOpen 7.11 (2018), pp. 845–849.

[11] Petra Jusková et al. “‘Basicles’: Microbial Growth and Production Monitoring in Giant Lipid Vesicles”. In: ACS Applied Materials & Interfaces 11.38 (2019), pp. 34698–34706. doi: 10.1021/acsami.9b12169.

[12] Pierre-Alain Monnard, Thomas Oberholzer, and PierLuigi Luisi. “Entrapment of nucleic acids in liposomes”. In: Biochimica et Biophysica Acta (BBA)-Biomembranes 1329.1 (1997), pp. 39–50.

[13] Lilian M Were et al. “Size, stability, and entrapment efficiency of phospholipid nanocapsules containing polypeptide antimicrobials”. In: Journal of agricultural and food chemistry 51.27 (2003), pp. 8073–8079.

[14] Marija Brgles et al. “Entrapment of ovalbumin into liposomes—factors affecting entrapment efficiency, liposome size, and zeta potential”. In: Journal of liposome research 18.3 (2008), pp. 235–248.

[15] Yongju He et al. “Influence of probe-sonication process on drug entrapment efficiency of liposomes loaded with a hydrophobic drug”. In: International Journal of polymeric materials and polymeric biomaterials 68.4 (2019), pp. 193–197.

[16] Benyamin Hoseini et al. “Optimizing nanoliposomal formulations: Assessing factors affecting entrapment efficiency of curcumin-loaded liposomes using machine learning”. In: International Journal of Pharmaceutics 646 (2023), p. 123414.

[17] Johan Hurtig and Owe Orwar. “Injection and transport of bacteria in nanotube–vesicle networks”. In: Soft matter 4.7 (2008), pp. 1515–1520.

[18] Lucas Le Nagard et al. “Encapsulated bacteria deform lipid vesicles into flagellated swimmers”. In: Proceedings of the National Academy of Sciences 119.34 (2022), e2206096119.

[19] Heidi MJ Weakly et al. “Several common methods of making vesicles (except an emulsion method) capture intended lipid ratios”. In: Biophysical journal 123.19 (2024), pp. 3452–3462.

[20] Kim S. Horger et al. “Films of Agarose Enable Rapid Formation of Giant Liposomes in Solutions of Physiologic Ionic Strength”. In: Journal of the American Chemical Society 131.5 (2009), pp. 1810–1819.

[21] Kim S Horger et al. “Hydrogel-assisted functional reconstitution of human P-glycoprotein (ABCB1) in giant liposomes”. In: Biochimica et Biophysica Acta (BBA)-Biomembranes 1848.2 (2015), pp. 643–653.

[22] A. D. Bangham, M. M. Standish, and J. C. Watkins. “Diffusion of Univalent Ions across the Lamellae of Swollen Phospholipids”. In: Journal of Molecular Biology 13.1 (1965), 238–IN27.

[23] Tadashi Matsunaga, Toshifumi Sakaguchi, and Fumihiko Tadakoro. “Magnetite formation by a magnetic bacterium capable of growing aerobically”. In: Applied Microbiology and Biotechnology 35 (1991), pp. 651– 655.

[24] Tadashi Matsunaga et al. “Complete genome sequence of the facultative anaerobic magnetotactic bacterium Magnetospirillum sp. strain AMB-1”. In: DNA research 12.3 (2005), pp. 157–166.

[25] Liu Yu et al. “Experimental determination of the propulsion matrix of the body of helical Magnetospirillum magneticum cells”. In: Physical Review E 106.3 (2022), p. 034407.

[26] CJ Pierce et al. “Tuning bacterial hydrodynamics with magnetic fields”. In: Physical Review E 95.6 (2017), p. 062612.

[27] Lucas Le Nagard et al. “Growing Magnetotactic Bacteria of the Genus Magnetospirillum: Strains MSR-1, AMB-1 and MS-1”. In: Journal of Visualized Experiments 140 (2018), e58536. doi: 10.3791/58536.

[28] R.K. Thauer Wolfe R.S. and N. Pfennig. ““A ‘Capillary Racetrack’ Method for Isolation of Magnetotactic Bacteria.”“. In: FEMS Microbiology Letters 45 (1 1987), pp. 31–35. doi: 10.1016/03781097(87)90039-5..

[29] Nicolas Färber and Christoph Westerhausen. “Broad lipid phase transitions in mammalian cell membranes measured by Laurdan fluorescence spectroscopy”. In: Biochimica et Biophysica Acta (BBA)-Biomembranes 1864.1 (2022), p. 183794.

[30] Susana A Sánchez et al. “Laurdan generalized polarization: from cuvette to microscope”. In: Modern research and educational topics in microscopy 2 (2007), pp. 1007–1014.

[31] Bacalum, Mihaela, Bogdan Zorilă, and Mihai Radu. “Fluorescence Spectra Decomposition by Asymmetric Functions: Laurdan Spectrum Revisited”. In: Analytical Biochemistry 440.2 (2013), pp. 123–129.

[32] Matthias Garten et al. “Reconstitution of a Transmembrane Protein, the Voltage-Gated Ion Channel, KvAP, into Giant Unilamellar Vesicles for Microscopy and Patch Clamp Studies”. In: Journal of Visualized Experiments 95 (2015), e52281. doi: 10.3791/52281.

[33] Phillip Kuhn et al. “A Facile Protocol for the Immobilisation of Vesicles, Virus Particles, Bacteria, and Yeast Cells”. In: Integrative Biology 4.12 (2012), pp. 1550– 1555.

[34] Rafael B. Lira et al. “Posing for a Picture: Vesicle Immobilization in Agarose Gel”. In: Scientific Reports 6.1 (2016), p. 25254.

[35] Juyang Huang, Jeffrey T Buboltz, and Gerald W Feigenson. “Maximum solubility of cholesterol in phosphatidylcholine and phosphatidylethanolamine bilayers”. In: Biochimica et Biophysica Acta (BBA) Biomembranes 1417.1 (1999), pp. 89–100.

[36] Hammad A Faizi et al. “Bending rigidity of charged lipid bilayer membranes”. In: Soft Matter 15.29 (2019), pp. 6006–6013.

[37] Mohammad Abu Sayem Karal et al. “A review on the measurement of the bending rigidity of lipid membranes”. In: Soft Matter 19.43 (2023), pp. 8285–8304.

[38] Yareni A Ayala et al. “Thermal fluctuations of the lipid membrane determine particle uptake into Giant Unilamellar Vesicles”. In: Nature Communications 14.1 (2023), p. 65.

[39] J Grant Burgess et al. “Evolutionary relationships among Magnetospirillum strains inferred from phylogenetic analysis of 16S rDNA sequences”. In: Journal of bacteriology 175.20 (1993), pp. 6689–6694.

[40] Christopher T Lefèvre et al. “Insight into the evolution of magnetotaxis in Magnetospirillum spp., based on mam gene phylogeny”. In: Applied and environmental microbiology 78.20 (2012), pp. 7238–7248.

[41] Christian Sohlenkamp and Otto Geiger. “Bacterial membrane lipids: diversity in structures and pathways”. In: FEMS microbiology reviews 40.1 (2016), pp. 133– 159.

[42] Anthony G Jay and James A Hamilton. “Disorder amidst membrane order: standardizing laurdan generalized polarization and membrane fluidity terms”. In: Journal of fluorescence 27.1 (2017), pp. 243–249.

[43] Faith M. Harris, Katrina B. Best, and John D. Bell. “Use of Laurdan Fluorescence Intensity and Polarization to Distinguish between Changes in Membrane Fluidity and Phospholipid Order”. In: Biochimica et Biophysica Acta (BBA) -Biomembranes 1565.1 (2002), pp. 123– 128. doi: 10.1016/S0005-2736(02)00514-X.

[44] Hanna Orlikowska-Rzeznik et al. “Laurdan discerns lipid membrane hydration and cholesterol content”. In: The Journal of Physical Chemistry B 127.15 (2023), pp. 3382–3391.

[45] Richard M Epand, Diana Bach, and Ellen Wachtel. “In vitro determination of the solubility limit of cholesterol in phospholipid bilayers”. In: Chemistry and Physics of Lipids 199 (2016), pp. 3–10.

[46] Susana A Sanchez, Maria A Tricerri, and Enrico Gratton. “Laurdan generalized polarization fluctuations measures membrane packing micro-heterogeneity in vivo”. In: Proceedings of the National Academy of Sciences 109.19 (2012), pp. 7314–7319.

[47] Denitsa Mitkova et al. “Bending rigidity of phosphatidylserine–containing lipid bilayers in acidic aqueous solutions”. In: Colloids and Surfaces A: Physicochemical and Engineering Aspects 460 (2014), pp. 71– 78.

[48] Ankur Gupta et al. “Different membrane order measurement techniques are not mutually consistent”. In: Biophysical Journal 122.6 (2023), pp. 964–972.

[49] Alexandros Giannopoulos-Dimitriou et al. “Liposome stability and integrity”. In: Liposomes in drug delivery. Elsevier, 2024, pp. 89–121.

[50] Yao Lu, Giulia Allegri, and Jurriaan Huskens. “Vesicle-Based Artificial Cells: Materials, Construction Methods and Applications”. In: Materials Horizons 9.3 (2022), pp. 892–907. doi: 10.1039/d1mh01431e.

