## Supplemental Figure 1, Table 1, Video 1 description for "Optimizing the membrane composition of immobilized giant vesicles for effective entrapment and observation of motile bacteria"

### Supporting information

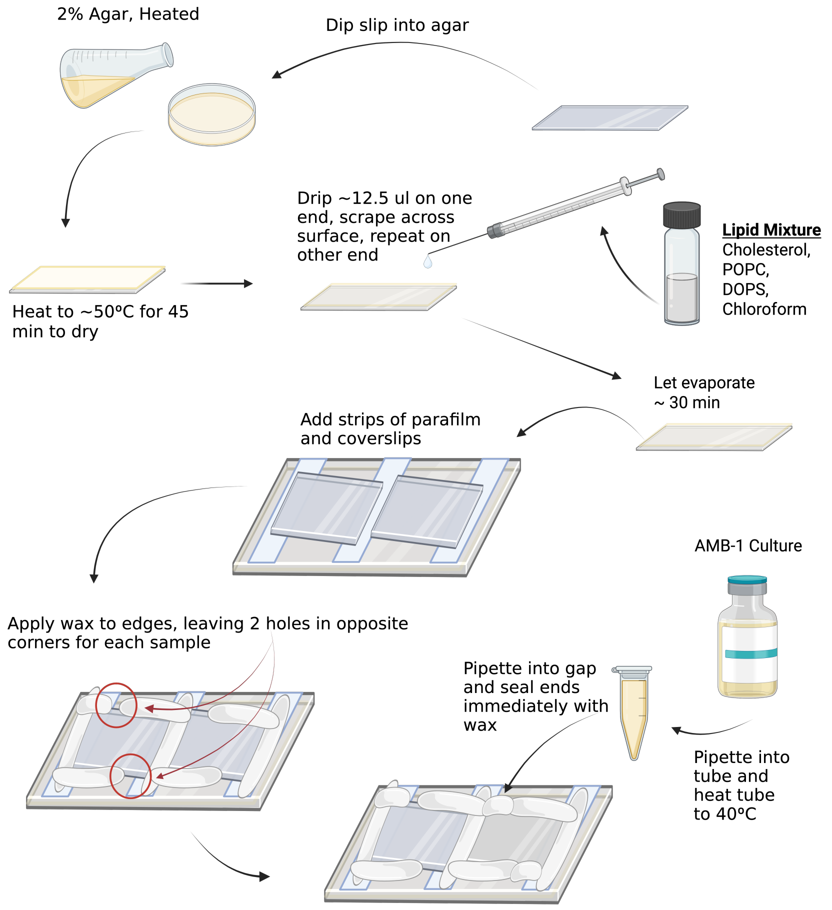


**S1 Figure. Sample Preparation.** Visual instructions for sample preparation of immobile GVs on agar using the swelling method. Created with BioRender.com/vu3zrr7.

**S1 Video. Entrapped Bacterium.** A video of a single bacterium entrapped in a GV. The bacterium was tracked using Trackmate (a plugin in ImageJ/Fiji) and an overlay of the bacterium shows the last 40 frames in which the bacterium traveled.

**S1 Table. Product/Ingredients Sources.**

| Product | Company | Catalog # | Lot |
|---|---|---|---|
| Agar | BioShop Canada | AGR003 | 1K63397 |
| Alexa Fluor 488 | Invitrogen | A10254 | 2160014 |
| Cholesterol (ovine) | Avanti Polar Lipids | 57-88-5 | 5092PPB110 |
| DPBS | Sigma-Aldrich | D8537 | - |
| DOPC | Avanti Polar Lipids | 840035C | 181PS-325 |
| Fe (III) Citrate | Sigma-Aldrich | F3388 | SLBR1615V |
| Fluorescent nanobeads | Invitrogen | F8803 | 2179346 |
| HEPES | BioShop Canada | HEP001 | 2A23376 |
| $KH_2PO_4$ | EM Science | PX1565-1 | 4233932 |
| Laurdan | Cayman Chemical | 19706 | 0600252-43 |
| $MgSO_4 \cdot 7H_2O$ | EM Science | MX0070-1 | 42140314 |
| $N_2$ | Air Liquide | A0492809 | - |
| $NaNO_3$ | Sigma-Aldrich | S5506 | MKBB6968 |
| NaOH | EMD Chemicals | SX0590-1 | 48311908 |
| POPC | Avanti Polar Lipids | 850457C | 5679CND216 |
| Potassium L-Lactate | Sigma-Aldrich | 60389 | BCBN6605V |
| Soy Bean Peptone | BD | 243620 | 5091836 |
| Trace Mineral Supplement | ATCC | MD-TMS | 61667728 |
| Well Plate (96) | Thermofisher Scientific | 165305 | - |
| Yeast Extract | Sigma-Aldrich | 70161 | 102634075 |
