## Supplementary figures and images for "Optimizing the membrane composition of immobilized giant vesicles for effective entrapment and observation of motile bacteria"

### Supplemental Figure 1

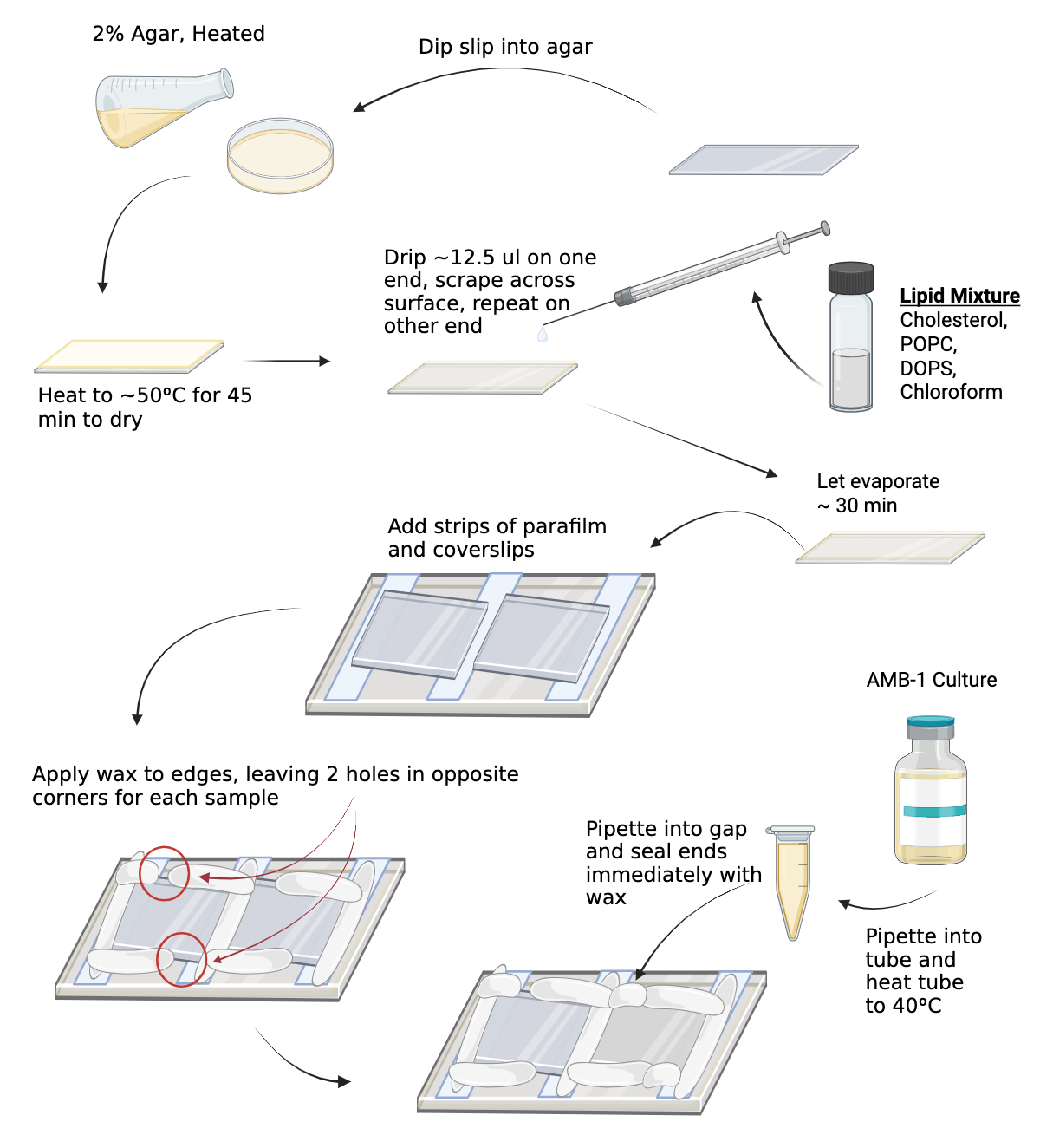
